# Collection of Sixteen High-Quality Human Head CAD Models Generated with SimNIBS 2.1 Using Connectome Subject Data within MATLAB^®^ Platform

**DOI:** 10.1101/480103

**Authors:** Aung Thu Htet, Gregory M. Noetscher, Edward H. Burnham, Aapo Nummenmaa, Sergey N. Makarov

## Abstract

The goal of this study is to introduce a collection of sixteen high-resolution, 2-manifold CAD compatible head models within the MATLAB platform available to all interested parties for electromagnetic and acoustic simulations. Each model contains skin, skull, CSF, GM, cerebellum, WM, and ventricles head compartments and possesses an “onion” topology: the grey matter shell is a container for white matter, ventricles, and cerebellum objects, the CSF shell contains the grey matter shell, the skull shell contains the CSF shell, and finally the skin or scalp shell contains the skull shell. The models are fully compatible with ANSYS ED FEM software, CST Studio Suite, Sim4Life/SEMCAD software, and other electromagnetic software packages.

The collection is based on MRI data from the Human Connectome Project (HCP) segmented using the SimNIBS 2.1/2.1.1 processing pipeline. The average number of triangular surface facets in a model is 866,000, the average triangle quality is 0.25, the average edge length is 1.48 mm, and the average surface mesh density or resolution is 0.57 points per mm^2^. If necessary, a finer model mesh can be created for every head using available MATLAB tools.

## 1. Introduction

For all three chief neurostimulation modalities – transcranial magnetic stimulation (TMS), transcranial electric stimulation (TES), and intracortical microstimulation (ICMS) – numerical computation of the electric fields within a patient-specific head model is the major and often only way to foster spatial targeting and obtain a quantitative measure of the required stimulation dose [1]. An important component of any simulation setup is the human head model itself, either a voxel model or a computer aided design (CAD) model represented by a set of solid objects [2].

Several open-source software packages currently exist that allow for automatic and semiautomatic generation of anatomically realistic CAD head models from individual MRI datasets. A comprehensive review can be found in [3]. Of specific note is the SimNIBS package [4]-[8] that uses FreeSurfer software [9],[10] for brain compartment segmentation (grey matter, white matter, cerebellum) and FSL [11],[12] for compatible skull/cerebrospinal fluid/ventricles segmentation when a combination of T1- and T2-weighted images is used as input.

At the same time, generic computational cranium models and collections of those continue to remain important for population-based studies as well as for testing accuracy, speed, and performance of various numerical algorithms and software packages. The most detailed anatomical CAD head model to date is arguably the MIDA model [13].

The Human Connectome Project (HCP) [14] provides a unique, open source, large-scale collection of about 1200 human head T1/T2 image datasets obtained with the isotropic resolution of 0.7 mm, along with the corresponding diffusion tensor imaging (DTL) data. Fifty such datasets have recently been segmented by E.G. Lee and colleagues [15],[16],[17] using SimNIBS, the well-known, open-source transcranial brain stimulation modeling software [4]-[8]. The resulting fifty CAD models, known as the Population Head Model Repository, are available from the website of the IT’IS Foundation, Switzerland [18].

The goal of this study is to provide a collection of sixteen high-resolution, 2-manifold, CAD ready head models within the MATLAB^®^ platform that are also fully compatible with ANSYS ED FEM software, CST Studio Suite, Sim4Life/SEMCAD software, and other electromagnetic software packages. Similar to the Population Head Model Repository [15],[16],[17] we also use the SimNIBS pipeline for automatic segmentation of structural T1 and T2 weighted MRI images of the Connectome database [14]. We extract the following major compartments: skin (or scalp), skull (outer surface), cerebrospinal fluid or CSF, which is simultaneously the inner surface of the skull, grey matter, cerebellum, white matter, and the ventricles. The entire model possesses an “onion” topology: the grey matter shell is a container for white matter, ventricles, and cerebellum objects, the CSF shell contains the grey matter shell, the skull shell contains the CSF shell, and finally the skin or scalp shell contains the skull shell.

In contrast to the Population Head Model Repository [15], [16], we did not perform any mesh postprocessing. Instead, we output the original data obtained with the most recent versions SimNIBS v. 2.1 and SimNIBS v. 2.1.1. In the Population Head Model Repository, this postprocessing performed with MeshMixer was focused on the skin (reducing mesh resolution) and the skull, where some models appeared to have very isolated divots that decreased the skull’s thickness, replacing it with more skin [17]. This postprocessing resulted in the appearance of non-manifold meshes which may be undesirable. Therefore, we independently segmented all fifty models of the Population Head Model Repository. After that, we have chosen sixteen models with a smooth skull surface. We also kept the high-resolution skin surface desirable for transcranial electric stimulation (TES) modeling and Electroencephalogram (EEG) modeling.

## 2. Materials and Methods

### 2.1 MRI Data

The MRI data were kindly provided by the HCP [14], WU-Minn Consortium (Principal Investigators: David Van Essen and Kamil Ugurbil; 1U54MH091657) funded by the 16 NIH Institutes and Centers that support the NIH Blueprint for Neuroscience Research; and by the McDonnell Center for Systems Neuroscience at Washington University. Users can sign up for a ConnectomeDB account and, after signing the Open Access Data Use Terms, can download more HCP data themselves as described on pp. 24-25 of the S1200 Reference Manual [21].

### 2.2 Segmentation

The T1 and T2 image files of the Connectome database [14] were used as input for the SimNIBS segmentation pipeline. For subject 101309, as an example, the command used was: `mri2mesh --all 101309 T1w.nii.gz T2w.nii.gz` where 101309 in this command is the unique Connectome’s subject ID. We updated from SimNIBS 2.1 to SimNIBS 2.1.1 at some time in the middle of the segmentation process. Subjects with IDs equal to or lower than 111716 were generated using SimNIBS 2.1 whereas subjects with IDs higher than that were segmented using SimNIBS 2.1.1.

Only the option `--all` was used. The option is “to run all reconstruction steps including volume meshing” according to `mri2mesh` documentation [19]. The segmentation time varies with each subject. Generally, segmentation of each head took from 10 to 12 hours using an Intel Xeon E5-2690 0 CPU 2.90GHz Linux server with 192 Gbytes of RAM.

The `mri2mesh` tool automatically generated the STL files for every individual head compartment. These files have been processed in MATLAB^®^ and in a commercial FEM electromagnetic software package ANSYS Electronics Desktop 2018.1.

### 2.3 Subject selection

After independently segmenting all fifty subjects used in the Population Head Repository [15]-[18], a visual inspection of the skull segmentation quality has been performed and the “acceptable” models have been retained. Figure 1 shows examples of acceptable and unacceptable outer skull shells.

**Figure 1.**
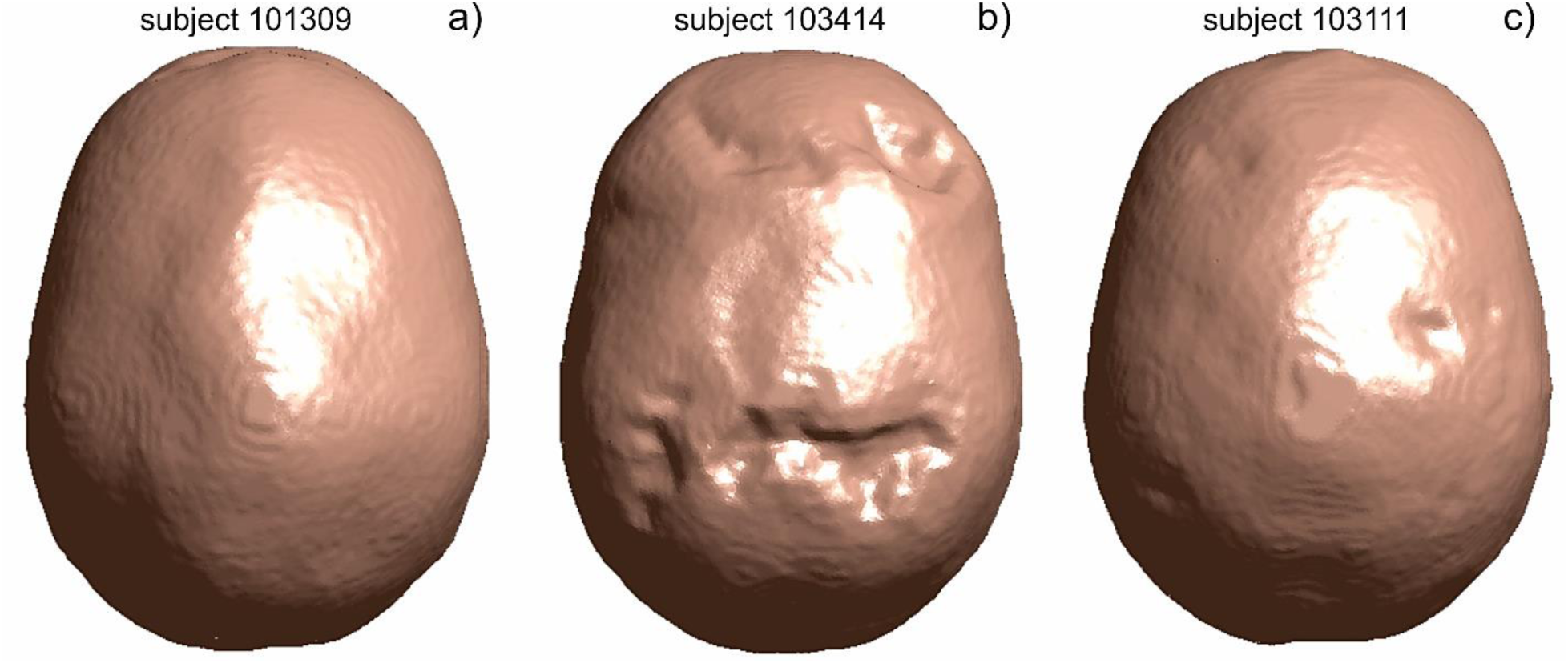
a) – Acceptable outer skin shell; b) – unacceptable outer skin shell; c) – unacceptable outer skin shell.

As a result, we ended up with sixteen acceptable models from the pool of fifty segmented models in total. We emphasize that an inspection of the CSF (or inner skull) surface and other compartments has also been performed.

### 2.4 Subject data

If restricted subject data (e.g. exact ages of subjects, health or substance use data, self-report psychiatric symptom data) is required, HCP needs to be informed and the Washington University IRB needs to be involved in the decision to allow or deny this.

## 3. Results

### 3.1 Data organization

Figure 2 shows the data organization chart. Every folder has the data for a specific head model: the folder name is the Connectome’s head ID. The data are given in common STL (STereoLithography) format and in MATLAB format (array of nodes P, array of facets *t*, and array of outer normal vectors *normals*).

**Figure 2.**
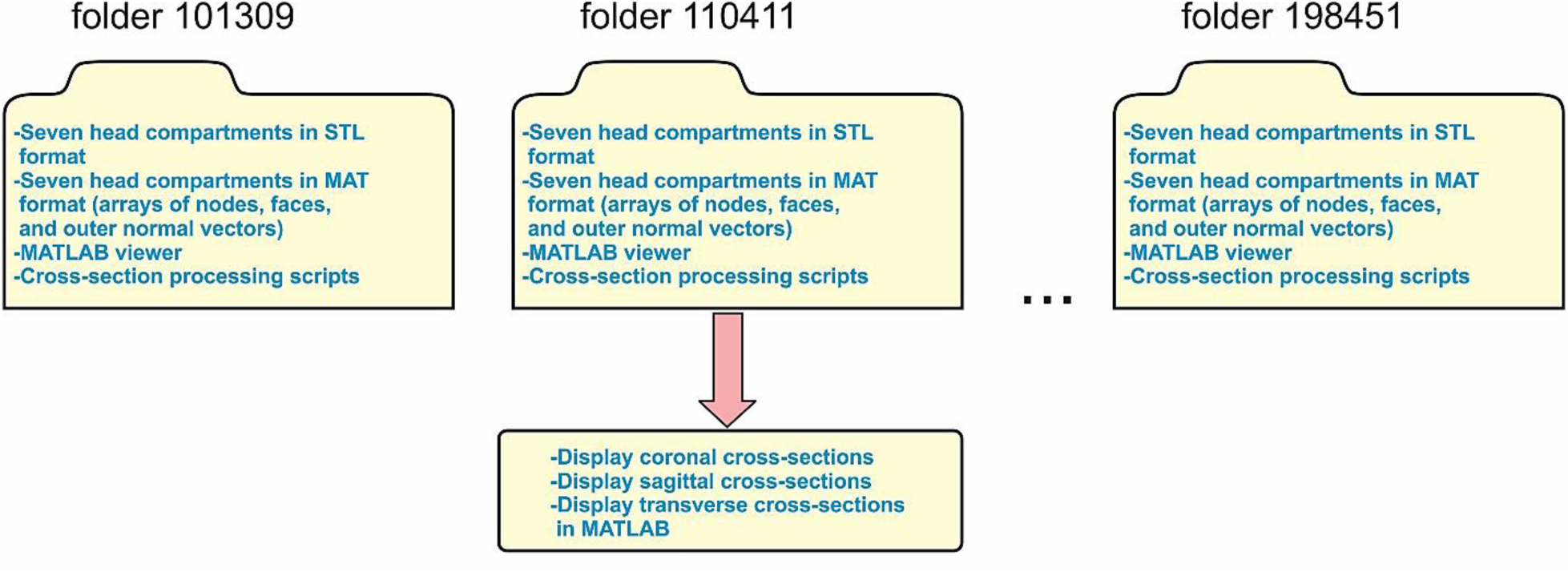
Data organization chart.

Every folder also has a number of simple MATLAB processing scripts which allow for a 3D visualization and model cross-section identification in any principal plane. Furthermore, the mesh characteristics can be computed as described below. As an example, Figure 3 shows three cross-section cuts for subject # 110411.

**Figure 3.**
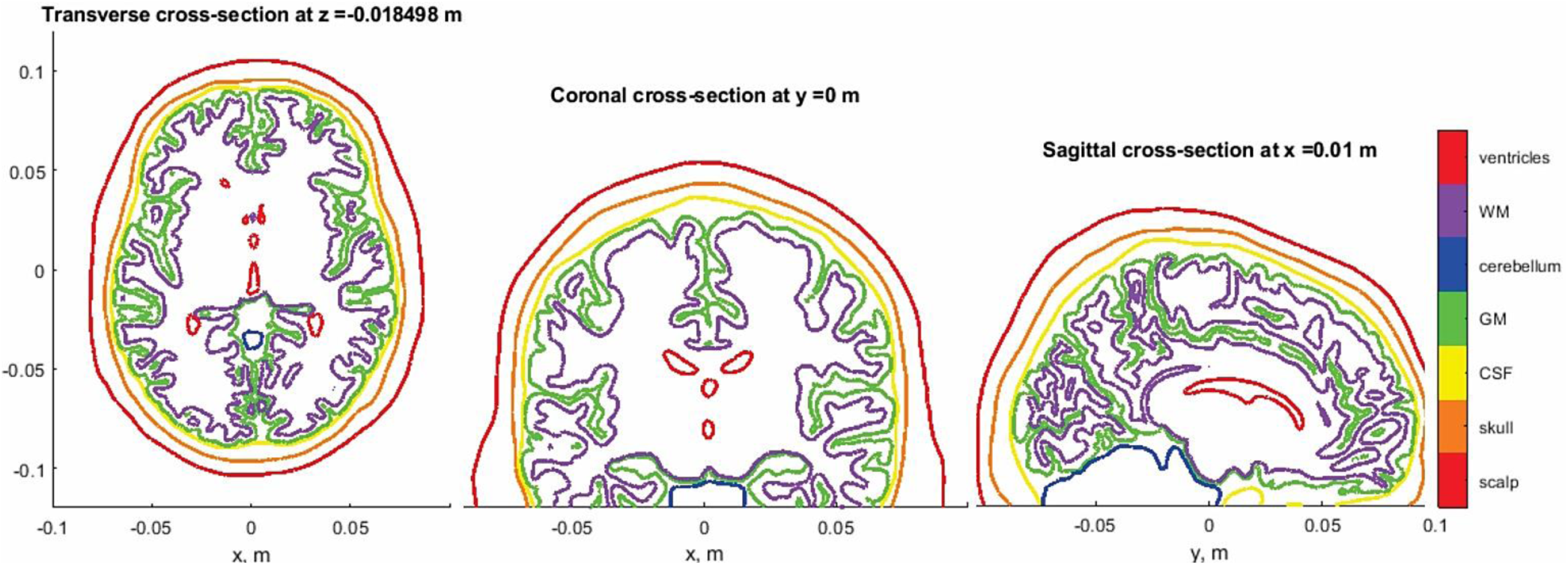
Three cross-section cuts for subject # 110411.

An outer folder has the STL to MAT conversion tools created by Dr. Pau Micó, Universitat Politècnica de València, Spain, and available from MATLAB Central [22]. Additionally, we also present a wrapper that collects and processes data from all sixteen head models.

### 3.2 Topological mesh characteristics

Table 1 reports the number of triangular faces in every head compartment of every dataset. The averaged total number of facets per head dataset is 866,000. Table 2 reports the mesh quality factor, *Q* [20], for every head compartment of every dataset. The average mesh quality over all compartments of all datasets is remarkably high; it is equal to 0.25. The skin and CSF (inner skull) shells possess the lowest mesh quality of the assembly. Table 3 reports the average edge length, *l*, mm of the triangular surface mesh for every head compartment of every dataset. The average edge length over all compartments of all datasets is 1.5 mm. Finally, Table 4 reports an average mesh resolution, *R*, in points/mm^2^ for every head compartment of every dataset. This parameter is in particular used by the developers of SimNIBS software. The average mesh resolution over all compartments of all datasets is 0.6 points per mm^2^.

**Table 1.** Number of triangular faces in each head compartment of every dataset.

| # | Head ID | Faces, skin | Faces, skull | Faces, CSF | Faces, GM | Faces, cerebel. | Faces, WM | Faces, ventricles | Faces, total |
| --- | --- | --- | --- | --- | --- | --- | --- | --- | --- |
| 1 | 101309 | 119990 | 119996 | 119994 | 222106 | 17136 | 235732 | 14958 | 849912 |
| 2 | 110411 | 119990 | 119992 | 119994 | 245886 | 17140 | 235608 | 14968 | 873578 |
| 3 | 117122 | 119992 | 119992 | 119994 | 240888 | 17152 | 235350 | 15004 | 868372 |
| 4 | 120111 | 119986 | 119992 | 119994 | 219610 | 17150 | 235120 | 12400 | 844252 |
| 5 | 122317 | 119986 | 119978 | 119990 | 245564 | 17142 | 235530 | 15004 | 873194 |
| 6 | 122620 | 119986 | 119988 | 119990 | 237744 | 17158 | 235420 | 15002 | 865288 |
| 7 | 124422 | 119990 | 119988 | 119992 | 251218 | 17146 | 235348 | 14998 | 878680 |
| 8 | 128632 | 119974 | 119990 | 119994 | 232810 | 17140 | 235346 | 14964 | 860218 |
| 9 | 130013 | 119990 | 119988 | 119996 | 238814 | 17142 | 235934 | 14994 | 866858 |
| 10 | 131722 | 119996 | 119986 | 119992 | 233072 | 17142 | 235844 | 14962 | 860994 |
| 11 | 138534 | 119992 | 119986 | 119988 | 239306 | 17146 | 235582 | 14960 | 866960 |
| 12 | 149337 | 119994 | 119990 | 119988 | 236482 | 17144 | 236078 | 14992 | 864668 |
| 13 | 149539 | 119994 | 119994 | 119994 | 246826 | 17142 | 235884 | 15004 | 874838 |
| 14 | 151627 | 119984 | 119988 | 119992 | 252570 | 17142 | 235250 | 14952 | 879878 |
| 15 | 160123 | 119900 | 119988 | 119994 | 241474 | 17142 | 235640 | 15008 | 869146 |
| 16 | 198451 | 119988 | 119988 | 119992 | 236556 | 17142 | 235992 | 14984 | 864642 |
| Avg |  | 119983 | 119989 | 119992 | 238808 | 17144 | 235604 | 14822 | 866342 |

**Table 2.**
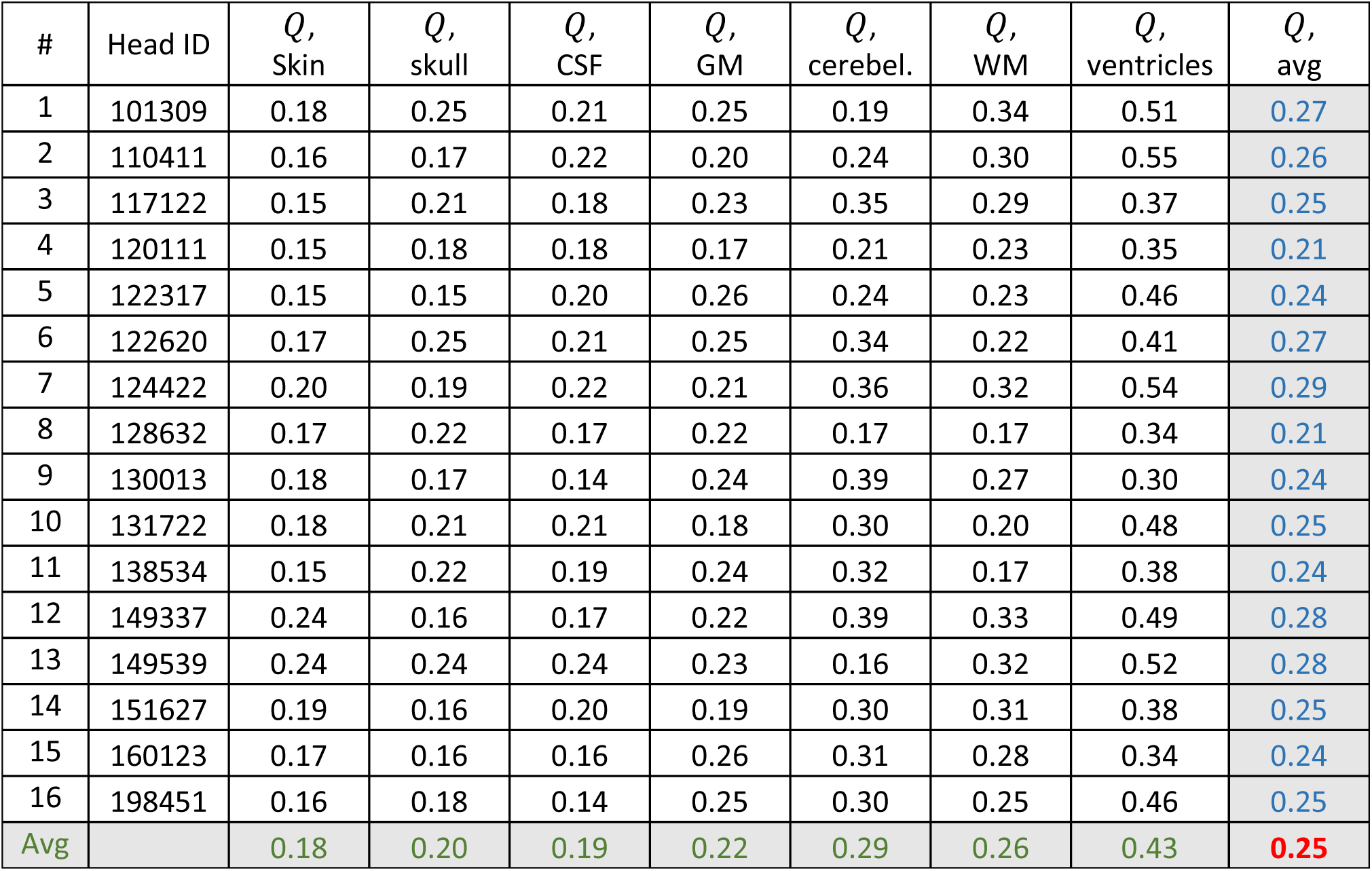
Mesh quality factor, *Q*, for each head compartment of every dataset.

**Table 3.** Average edge length, *l*, mm for each head compartment of every dataset.

| # | Head ID | $l$,<br>Skin | $l$,<br>skull | $l$,<br>CSF | $l$,<br>GM | $l$,<br>cerebel. | $l$,<br>WM | $l$,<br>ventricles | $l$,<br>avg |
| --- | --- | --- | --- | --- | --- | --- | --- | --- | --- |
| 1 | 101309 | 1.65 | 1.41 | 1.31 | 1.54 | 1.84 | 1.51 | 1.10 | 1.48 |
| 2 | 110411 | 1.61 | 1.40 | 1.30 | 1.49 | 1.85 | 1.46 | 1.18 | 1.47 |
| 3 | 117122 | 1.60 | 1.39 | 1.31 | 1.50 | 1.85 | 1.45 | 1.22 | 1.48 |
| 4 | 120111 | 1.59 | 1.39 | 1.31 | 1.52 | 1.81 | 1.46 | 1.12 | 1.46 |
| 5 | 122317 | 1.59 | 1.39 | 1.31 | 1.50 | 1.82 | 1.45 | 1.19 | 1.46 |
| 6 | 122620 | 1.63 | 1.41 | 1.31 | 1.52 | 1.87 | 1.45 | 1.24 | 1.49 |
| 7 | 124422 | 1.68 | 1.39 | 1.31 | 1.52 | 1.85 | 1.46 | 1.25 | 1.49 |
| 8 | 128632 | 1.64 | 1.40 | 1.31 | 1.54 | 1.85 | 1.48 | 1.19 | 1.49 |
| 9 | 130013 | 1.58 | 1.38 | 1.30 | 1.55 | 1.87 | 1.49 | 1.24 | 1.49 |
| 10 | 131722 | 1.64 | 1.40 | 1.31 | 1.53 | 1.85 | 1.50 | 1.18 | 1.49 |
| 11 | 138534 | 1.69 | 1.42 | 1.30 | 1.51 | 1.85 | 1.46 | 1.16 | 1.48 |
| 12 | 149337 | 1.64 | 1.39 | 1.30 | 1.54 | 1.87 | 1.48 | 1.23 | 1.49 |
| 13 | 149539 | 1.65 | 1.38 | 1.31 | 1.50 | 1.85 | 1.46 | 1.20 | 1.48 |
| 14 | 151627 | 1.66 | 1.39 | 1.31 | 1.48 | 1.86 | 1.42 | 1.23 | 1.48 |
| 15 | 160123 | 1.68 | 1.39 | 1.30 | 1.48 | 1.83 | 1.45 | 1.18 | 1.47 |
| 16 | 198451 | 1.63 | 1.39 | 1.31 | 1.57 | 1.85 | 1.52 | 1.19 | 1.49 |
| Avg |  | 1.64 | 1.39 | 1.31 | 1.52 | 1.85 | 1.47 | 1.19 | 1.48 |

**Table 4.** Average mesh resolution, *R*, points/mm^2^ for each head compartment of every dataset.

| # | Head ID | $R$,<br>skin | $R$,<br>skull | $R$,<br>CSF | $R$,<br>GM | $R$,<br>cerebel. | $R$,<br>WM | $R$,<br>ventricles | $R$,<br>avg |
| --- | --- | --- | --- | --- | --- | --- | --- | --- | --- |
| 1 | 101309 | 0.44 | 0.61 | 0.70 | 0.50 | 0.35 | 0.52 | 0.98 | 0.59 |
| 2 | 110411 | 0.46 | 0.61 | 0.71 | 0.53 | 0.35 | 0.56 | 0.86 | 0.58 |
| 3 | 117122 | 0.47 | 0.62 | 0.70 | 0.53 | 0.35 | 0.56 | 0.79 | 0.57 |
| 4 | 120111 | 0.47 | 0.62 | 0.70 | 0.51 | 0.37 | 0.56 | 0.95 | 0.60 |
| 5 | 122317 | 0.47 | 0.62 | 0.70 | 0.53 | 0.36 | 0.57 | 0.84 | 0.58 |
| 6 | 122620 | 0.45 | 0.61 | 0.69 | 0.51 | 0.34 | 0.57 | 0.77 | 0.56 |
| 7 | 124422 | 0.43 | 0.62 | 0.70 | 0.51 | 0.35 | 0.56 | 0.76 | 0.56 |
| 8 | 128632 | 0.45 | 0.62 | 0.70 | 0.50 | 0.35 | 0.55 | 0.84 | 0.57 |
| 9 | 130013 | 0.48 | 0.63 | 0.71 | 0.49 | 0.34 | 0.53 | 0.78 | 0.57 |
| 10 | 131722 | 0.45 | 0.61 | 0.70 | 0.51 | 0.35 | 0.53 | 0.86 | 0.57 |
| 11 | 138534 | 0.42 | 0.59 | 0.71 | 0.52 | 0.35 | 0.56 | 0.88 | 0.58 |
| 12 | 149337 | 0.45 | 0.62 | 0.71 | 0.50 | 0.34 | 0.54 | 0.78 | 0.56 |
| 13 | 149539 | 0.44 | 0.63 | 0.70 | 0.52 | 0.35 | 0.56 | 0.82 | 0.57 |
| 14 | 151627 | 0.43 | 0.62 | 0.70 | 0.54 | 0.35 | 0.59 | 0.78 | 0.57 |
| 15 | 160123 | 0.42 | 0.62 | 0.70 | 0.54 | 0.36 | 0.56 | 0.85 | 0.58 |
| 16 | 198451 | 0.45 | 0.62 | 0.70 | 0.48 | 0.35 | 0.52 | 0.83 | 0.57 |
| Avg |  | 0.45 | 0.62 | 0.70 | 0.51 | 0.35 | 0.55 | 0.84 | 0.57 |

### 3.3 Specific conditions for CAD models

The following two conditions are required for a true CAD human body model [20]: (1) a 3D triangular mesh representing a solid object must not have holes; (2) the surface of a well-behaved triangular mesh in 3D must satisfy the so-called manifold condition. A mesh is 2-manifold if every node of the mesh has a disk-shaped neighborhood of triangles. This neighborhood can be continuously deformed to an open disk. Every edge of a 2-manifold mesh is a manifold edge with only two attached triangles. All other meshes are non-manifold meshes and are not suitable for FEM analysis. Figure 4 gives examples of a manifold edge, a non-manifold edge and a non-manifold mesh with a non-manifold node.

**Figure 4.**
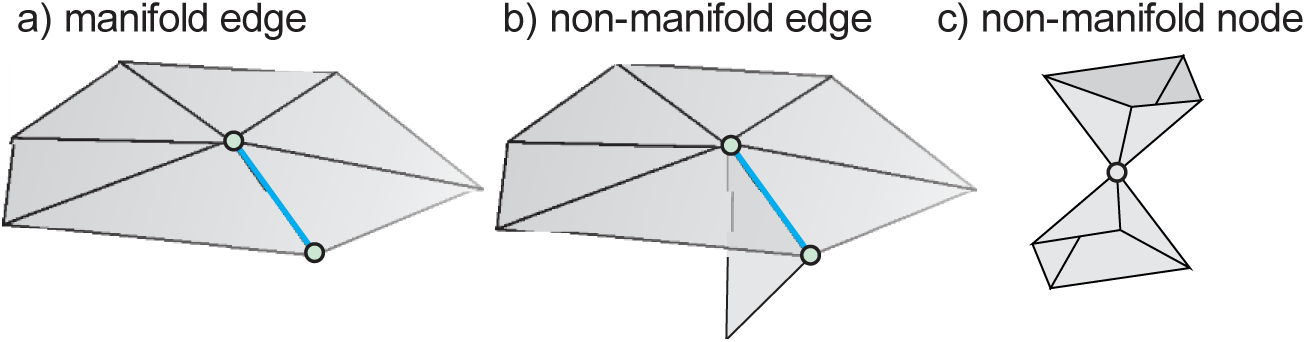
a) – Examples of a manifold edge; b) – non-manifold edge, and c) – non-manifold node.

In order to check the CAD conditions, we applied the mesh checker of the commercial FEM software ANSYS Electronics Desktop (ANSYS Maxwell) 2018.1 with the check option “strict”. Each head compartment of every dataset has been checked by creating a separate ANSYS project and importing the corresponding STL data. The results showed that all the surface meshes passed the ANSYS check without errors.

### 3.4 Comparison with the Population Head Repository

As compared to the Population Head Repository [18], we have nearly doubled the resolution and significantly increased triangle quality for the skin surface. Also, all present CAD meshes are 2-manifold, which is not the case for the Population Head Repository.

## 4. Conclusions

In this paper, we have described a collection of sixteen high-resolution, 2-manifold, CAD compatible head models within the MATLAB^®^ platform available to all interested parties for electromagnetic and acoustic simulations. Each model contains skin, skull, CSF, GM, cerebellum, WM, and ventricles head compartments and possesses an “onion” topology: the grey matter shell is a container for white matter, ventricles, and cerebellum objects, the CSF shell contains the grey matter shell, the skull shell contains the CSF shell, and finally the skin or scalp shell contains the skull shell. The models are fully compatible with ANSYS ED FEM software, CST Studio Suite, Sim4Life/SEMCAD software, and other electromagnetic software packages.

The collection is based on MRI data from the Human Connectome Project segmented using the SimNIBS 2.1/2.1.1 processing pipeline. The average number of triangular surface facets in a model is 866,000, the average triangle quality is 0.25, the average edge length is 1.48 mm, and the average surface mesh density or resolution is 0.57 points per mm^2^.

The collection is available via MATLAB Central at https://www.mathworks.com/matlabcentral/fileexchange/69517-collection-of-sixteen-high-quality-human-head-cad-models.

## Acknowledgements

The authors wish to thank Dr. Guilherme Saturnino of Danish Research Centre for Magnetic Resonance (SimNIBS), Mr. Erik Lee of Northwestern University, and Dr. Jennifer Elam of Human Connectome Project, Washington University School of Medicine for help with this project. The authors are also thankful to Sim4Life staff (Zurich, Switzerland) for providing their electromagnetic simulation software.

